# Population genomics of *Anopheles darlingi*, the principal South American malaria vector mosquito

**DOI:** 10.1101/2025.03.13.643102

**Authors:** Jacob A. Tennessen, Raphael Brosula, Estelle Chabanol, Sara Bickersmith, Angela M. Early, Margaret Laws, Katrina A. Kelley, Maria Eugenia Grillet, Dionicia Gamboa, Eric R. Lucas, Jean-Bernard Duchemin, Martha L. Quiñones, Maria Anice Mureb Sallum, Eduardo S. Bergo, Jorge E. Moreno, Sanjay Nagi, Nicholas J. Arisco, Mohini Sooklall, Reza Niles-Robin, Marcia C. Castro, Horace Cox, Mathilde Gendrin, Jan E. Conn, Daniel E. Neafsey

**Affiliations:** Harvard T.H. Chan School of Public Health; Boston, MA USA; Broad Institute; Cambridge, MA USA; Institut Pasteur de la Guyane; Cayenne, French Guiana; New York State Department of Health, Wadsworth Center; Albany, NY USA; Instituto de Zoología y Ecología Tropical, Facultad de Ciencias, Universidad Central de Venezuela; Caracas, Venezuela; Laboratorio de Malaria: Parásitos y Vectores, Laboratorios de Investigación y Desarrollo, Facultad de Ciencias e Ingeniería, Universidad Peruana Cayetano Heredia; Lima, Peru; Liverpool School of Tropical Medicine; Liverpool, UK; Universidad Nacional de Colombia; Bogotá, Colombia; Faculdade de Saúde Pública, Universidade de São Paulo; São Paulo, Brazil; Instituto Pasteur de São Paulo; São Paulo, Brazil; Instituto de Altos Estudios Dr. Arnoldo Gabaldón, Centro de Investigaciones de Campo Francesco Vitanza; Bolivar, Venezuela; Vector Control Services, Ministry of Health; Georgetown, Guyana; Department of Biomedical Sciences, College of Integrated Health Sciences, State University of New York at Albany; Albany, NY USA

## Abstract

Malaria in South America remains a serious public health problem. *Anopheles* (*Nyssorhynchus*) *darlingi* is the most important malaria vector across tropical Latin America. Vector-targeted disease control efforts require a thorough understanding of mosquito demographic and evolutionary patterns. We present and analyze whole genomes of 1094 *A. darlingi* (median depth 18x) from six South American countries. We observe deep geographic population structure, high genetic diversity including thirteen putative segregating inversions, and no evidence for cryptic sympatric taxa despite high interpopulation divergence. Strong signals of selection are plausibly driven by insecticides, especially on cytochrome P450 genes, one of which we validated experimentally. Our results will facilitate effective mosquito surveillance and control, while highlighting ongoing challenges that a diverse vector poses for malaria elimination in the western hemisphere.

---

There are approximately half a million malaria cases per year in the Americas (*1*), and the disease remains a stubbornly persistent health crisis in this region. Some locations have recently endured rapid upswings in case numbers (*1, 2*), such that case counts for the Americas overall have declined little over the past decade (*1*), dousing optimism for imminent malaria elimination in some countries. Furthermore, South America has been a hotspot for drug resistance emergence in the *Plasmodium* spp. parasites that cause malaria (*3–5*), threatening control efforts around the world. Successful elimination of neotropical malaria will require a greater understanding of its transmission determinants, including the population biology and adaptations of its most widespread and diverse mosquito vector, *Anopheles* (*Nyssorhynchus*) *darlingi*. This species is the primary vector throughout the Guiana Shield and Amazon Basin and an important vector across much of the rest of tropical Latin America, efficiently transmitting both *Plasmodium vivax* and *Plasmodium falciparum* (*6, 7*).

There is deep evolutionary divergence between malaria vectors in the neotropics relative to the rest of the world. The *Nyssorhynchus* subgenus, which contains *A. darlingi*, is confined to the neotropics (*8*), and is arguably a distinct genus (*9*). It diverged from the *Cellia* subgenus, which contains the principal vector of malaria in Africa (*Anopheles gambiae*), approximately 90 million years ago (*10*). Despite extensive population genomic analysis of malaria vectors in Africa and Asia (*11–14*), comparable work on neotropical anophelines has been restricted to relatively small population genetic datasets. In particular, it has remained unclear whether *A. darlingi* is an assemblage of cryptic, morphologically-identical species, as genomic datasets have recently elucidated for the *A. gambiae* complex (*15*), *A. minimus* complex (*13*), and *A. funestus* complex (*16*). Population genetic subdivisions have been observed in *A. darlingi* on either side of the Andes (*17*) and between the Amazon basin and southern Brazil (*18, 19*), however no genomic investigation has been conducted to determine whether sympatric cryptic species exist.

Sympatric cryptic species, first detected in African anophelines through partial sterility in laboratory crossing experiments, have complicated efforts to accurately measure insecticide resistance phenotypes and identify genotypic markers of insecticide resistance in African vectors since the advent of large-scale insecticidal control almost a century ago (*20, 21*). While other *Anopheles* species repeatedly evolve insecticide resistance through target site polymorphisms as well as metabolically-acting factors like cytochrome P450s (*22–24*), in *A. darlingi* insecticide resistance alleles have not been detected (*25*) and observations of phenotypic resistance have been rare and restricted to certain geographic regions and classes of insecticide (*26–28*).

Here we present a population genomic analysis of *A. darlingi*, an animal with one of the greatest impacts on public health in the Americas. In addition to describing overall patterns of genetic diversity, a major goal of this study has been to assess whether this species exhibits the characteristic evolutionary patterns of African and Asian *Anopheles* that have been particularly vexing to vector control goals, including partially interbreeding cryptic taxa, strikingly high nucleotide diversity partitioned into large segregating haplotype blocks, and insecticide resistance via target-site and metabolic mechanisms.

## Pronounced geographical population structure without sympatric cryptic taxa

We generated whole-genome sequences for 1094 adult female samples of *A. darlingi* from 16 locations in six South American countries (Fig. 1A; tables S1, and S2). Our extensive sampling represents most of the species’ range with some exceptions (e.g. Central America and much of southern Brazil). Excluding eight samples with elevated heterozygosity indicative of contamination (fig. S1), and 62 samples with median coverage ≤1x, there were 1024 usable samples (median coverage 20x). Collection localities were ecologically diverse (Fig. 1B). We detected DNA from multiple vertebrate species including humans and domesticated animals, presumably deriving from recent blood meals, as well as DNA from *P. falciparum* and *P. vivax*, which was significantly more common within samples positive for human DNA (P < 0.01; table S2). Rates of disease transmission will depend on how often mosquitoes bite humans relative to other vertebrates, in turn influenced by ecological factors and human behavior.

**Fig. 1.**
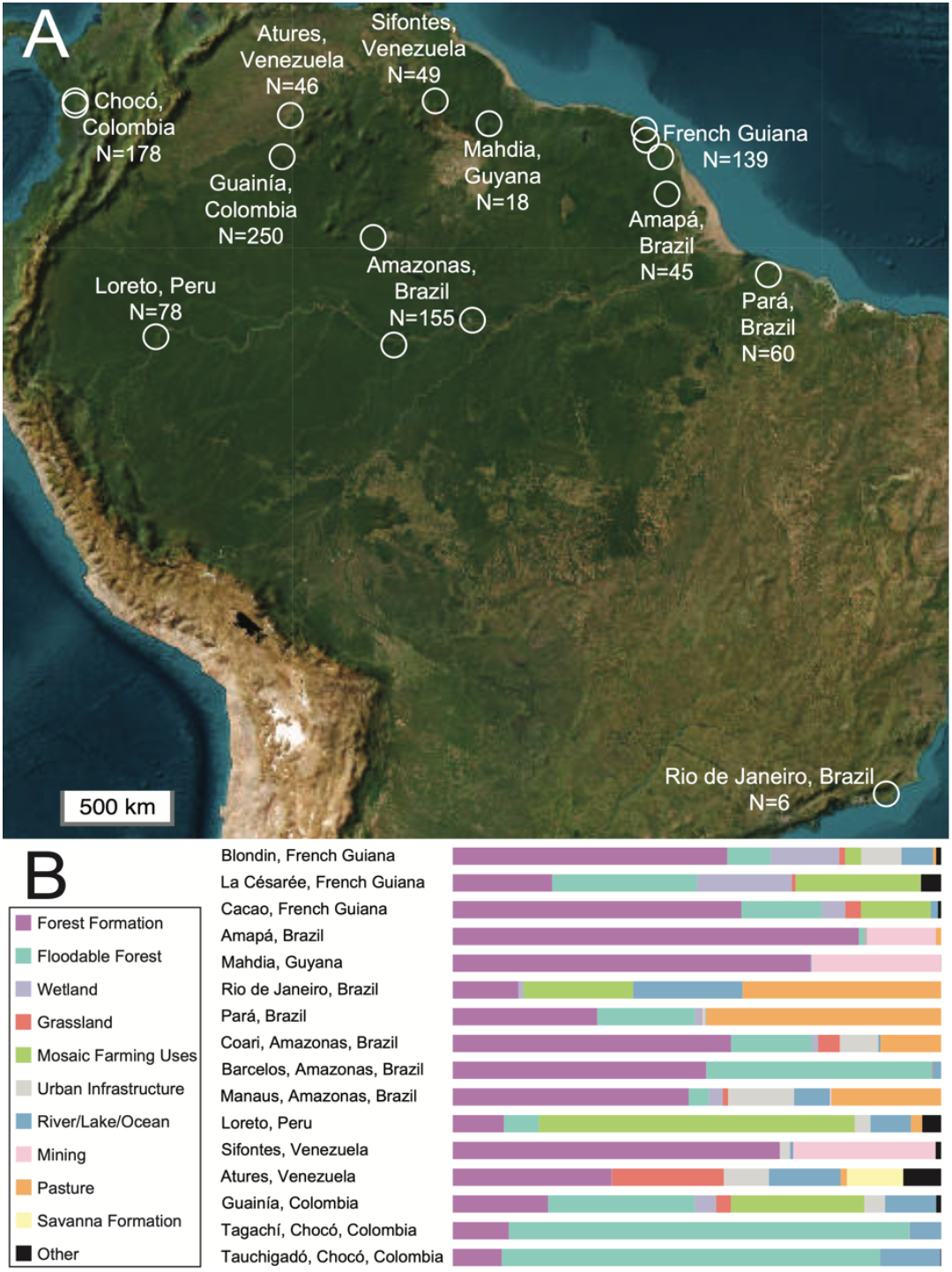
Collection sites. **(A)** Collection localities with counts of usable samples in six countries. **(B)** Proportion of land cover and land use type in the surrounding 5 km area for all collection sites.

We observe genetic divergence across populations, as seen in other widespread South American insects (*29–32*), confirming mitochondrial observations (*33*). Ancestry analysis with ADMIXTURE predicted eight ancestral populations, closely corresponding to contemporary collection sites, with little evidence of admixture in most locales (Fig. 2A). Ancestry models with fewer than eight populations are suboptimal but indicate that the most fundamental splits (K = 2 or 3) are among an Amazon/Atlantic clade (French Guiana, Amapá, Guyana, Rio de Janeiro, Pará, Amazonas, Peru), an Orinoco clade (Sifontes, Atures, Guainía), and a Chocó clade west of the Andes (fig. S2). Principal component analysis also clusters samples in a manner consistent with geography, with the largest split separating Colombian and Venezuela mosquitoes from all others (Fig. 2B). Similar results are obtained for genomic partitions such as chromosome arms (fig. S3) and for geographical subsets of samples (fig. S4). A population phylogeny with TreeMix is consistent with these results: first the Chocó clade branches off, followed by Guainía/Venezuela (Orinoco clade), leaving an Amazon/Atlantic clade within which Amazonas/Peru splits from the remaining eastern populations (Fig. 2C). The Andes represent an intuitive and frequently documented barrier to gene flow in many regional species (*31*), consistent with the isolation of Chocó *A. darlingi* (*17*). More surprising is the distinction between the two main eastern clades based around drainage basins that are not separated by obvious physical barriers preventing gene flow: Orinoco versus Amazon/Atlantic. These clades split Sifontes, Venezuela from Mahdia, Guyana, separated by only 270 km of low-elevation suitable habitat.

**Fig. 2.**
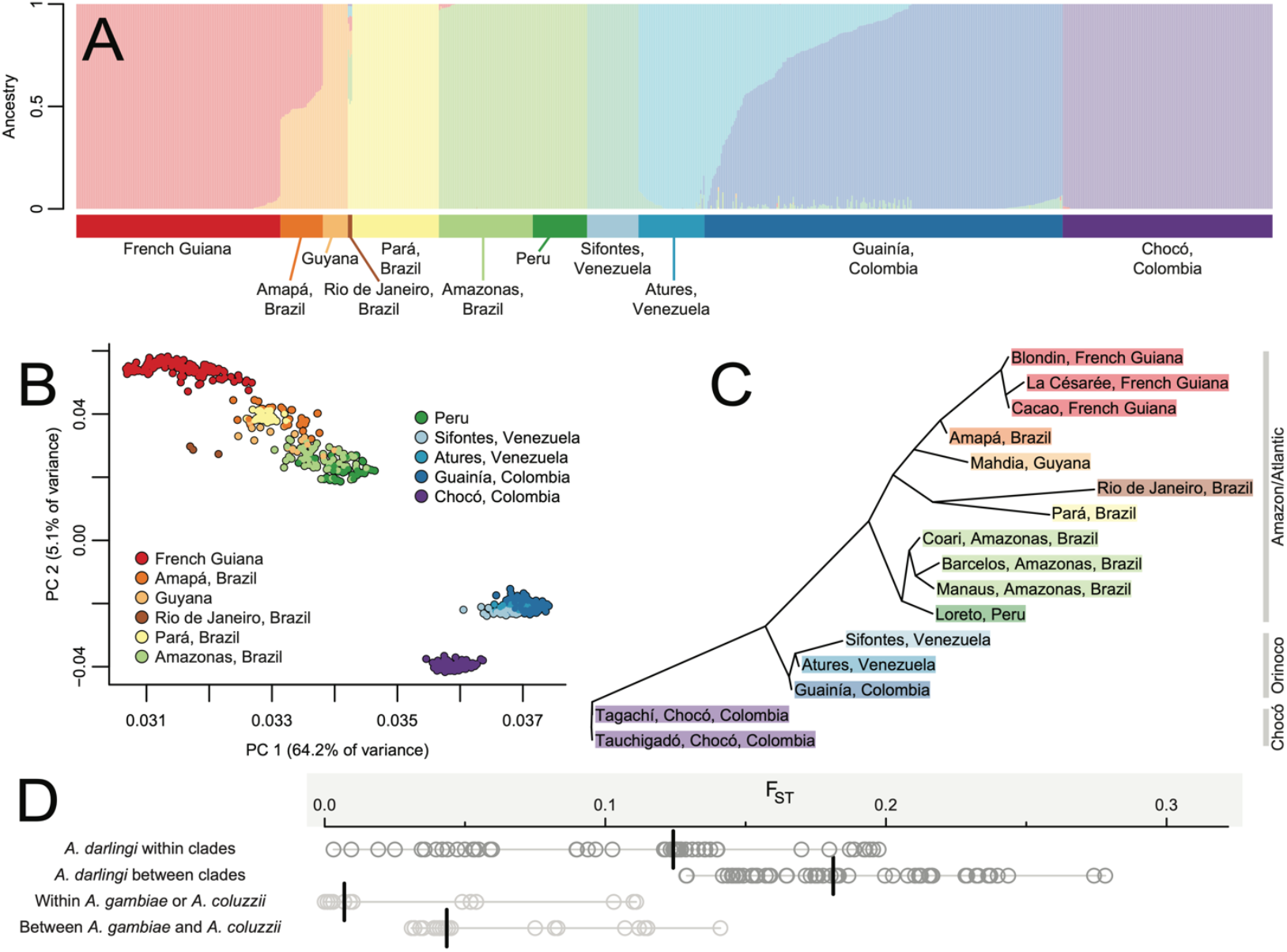
Population structure. **(A)** Ancestry assignment via ADMIXTURE identifies eight ancestral populations, closely corresponding to the contemporary populations of French Guiana, Guyana, Pará, Amazonas/Peru, Sifontes, Atures, Guainía, and Chocó. Only Amapá, Rio de Janeiro, and Guainía show substantial admixture. **(B)** Principal component analysis separates samples geographically. **(C)** Sampling site phylogeny generated with TreeMix, showing three major clades: Amazon/Atlantic, Orinoco, and Chocó. **(D)** Pairwise F_ST_ among fifteen *A. darlingi* collection sites (Rio de Janeiro excluded for sample size), classified as within major clades (from part C) (e.g. Guyana vs. Peru) or between clades (e.g. Guyana vs. Sifontes). Within- and between-species values for African *A. gambiae* and *A. coluzzii* shown for comparison. For all four sets of points across both continents, median geographic distance is similarly large (1000-2000 km). Black vertical lines show medians.

There is no evidence of sympatric cryptic taxa: samples from the same site are always genetically homogeneous with closely shared ancestry (Fig. 2A,B) and do not depart systematically from Hardy-Weinberg equilibrium (fig. S5). However, further study will be needed to determine whether geographically distinct clades show any reproductive isolation.

Pairwise F_ST_ is high, especially between the major clades (median 0.18), exceeding F_ST_ values observed in *A. gambiae* and *A. coluzzii* (*11, 12*) (Fig. 2D, fig. S6), in which relatively low divergence (F_ST_ < 0.01) is typical between continental populations separated by vast physical distance up to 4500 km. F_ST_ between the *A. darlingi* major clades even surpasses F_ST_ between African species (< 0.15). While *A. gambiae* relies on high-altitude wind dispersal to facilitate gene flow (*34*), it is unknown whether *A. darlingi* can utilize a similar mechanism. Plausibly, long-range migration has been more adaptive in Africa than in South America due to increased aridity and seasonality of wetlands, leading to the distinct population genetic patterns on each continent.

The deep population structure we observe across South America has both positive and negative consequences for mosquito-focused interventions. Mitochondrial DNA has suggested the Amazon River as a barrier to gene flow in *A. darlingi* (*18*), but we observe relatively high genetic affinity between Amapá and Pará on opposite sides of the Amazon Delta (Figs. 1 and 2), similar to the pattern seen in *A. aquasalis* across the same range (*35*), and mirroring the close kinship between *A. darlingi* populations separated by the Orinoco. Instead of rivers as barriers, we observe the opposite: boundaries between watersheds demarcate the major genetic divisions. Gene flow may be facilitated by waterways due to the dependence of *A. darlingi* on aquatic breeding sites. Interpopulation connectivity may also be enhanced by anthropogenic activity along rivers including deforestation leading to favorable disturbed habitat (*32, 36*) or transport in vehicles or boats. If a population were to be extirpated as reported in Suriname (*37*), there could be a long delay before the habitat is recolonized via migration from neighboring countries, unless enabled by human activity.

Limited gene flow also indicates that a new drug resistance allele will not spread quickly beyond its population of origin. While we are unaware of plans to modify *A. darlingi* with gene drives as is being attempted on other species (*38*), our results indicate that a future gene drive implementation in these mosquitoes would likely require repeated launches in several geographic locations, potentially requiring regional coordination.

## High genetic diversity partitioned into large haplotype blocks

Genetic diversity is extremely high, with heterozygosity >2% in most samples (Fig. 3A). Synonymous-site nucleotide diversity (π_S_) is 3.2% on autosomes, comparable to the famously high nucleotide diversity of the *A. gambiae* complex (*11*), and 2.3% on the X (Fig. 3B), 2-fold higher than *A. gambiae*. X chromosome π is approximately 75% of autosomal π, reflecting the expected X/autosome effective population size ratio and suggesting that X diversity in *A. darlingi* is not impacted by the factors that reduce it further in *A. gambiae*, such as sex ratio skew.

**Fig. 3.**
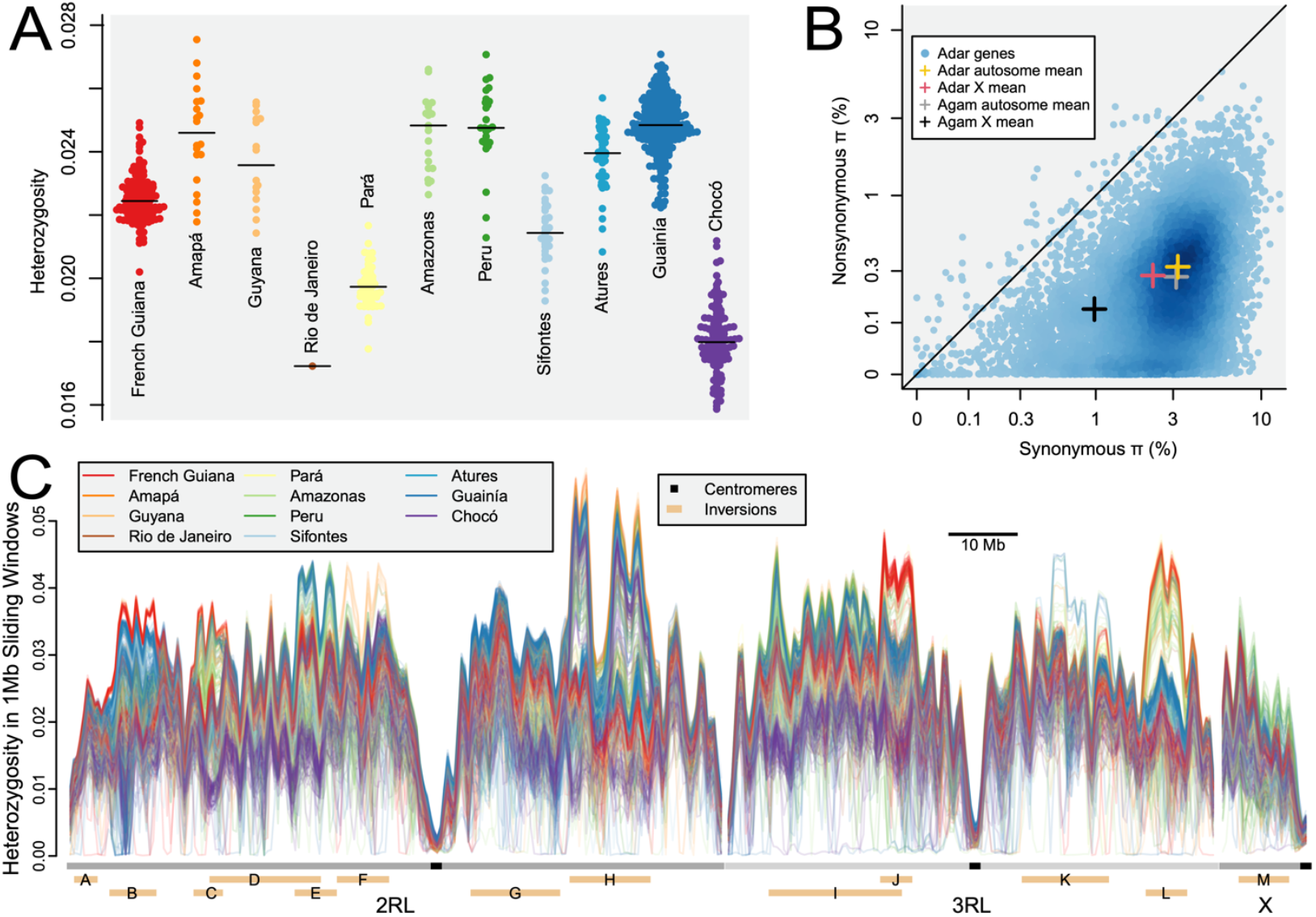
Genetic diversity. **(A)** Median heterozygosity per sample, arranged by population. **(B)** Synonymous and nonsynonymous nucleotide diversity (π) per gene. Dotted lines = mean values. Solid line = π_N_/π_S_ of 1. Crosses show mean values for X and autosomal genes, both for *A. darlingi* (Adar) and for the *A. gambiae* complex (Agam) as a comparison. **(C)** Heterozygosity per sample in sliding 1 Mb windows. Samples colored by population. Tan rectangles (bottom) indicate inversions, designated by capital letters. Peaks of elevated heterozygosity in some individuals typically overlap inversions and represent heterozygotes for the two haplotype lineages, with heterozygosity often up to two-fold higher than homozygotes. Occasionally, otherwise diverse individuals have regions of near-zero heterozygosity for over a megabase, indicating homozygosity along a long haplotype.

Tajima’s D in *A. darlingi* is positive and relatively close to zero (median per population 0.26, fig. S7), contrasting with the very negatively-skewed site frequency spectra of *A. gambiae* (Tajima’s D < -1.5) attributable to explosive population growth facilitated by human-modified habitats (*11*). All *A. darlingi* clades show comparable genetic diversity and site frequency spectra, suggesting relative stability in long-term population size and persistence, with potentially slight decreases in abundance, especially in relatively low diversity populations like Chocó, Sifontes, and Pará (Fig. 3A). Thus, deforestation, urbanization, and other anthropogenic changes across Latin America have not resulted in an exponential increase in *A. darlingi* as similar changes have for African mosquitoes, likely because *A. darlingi* is less anthropophilic (*39, 40*). However, very recent expansions in response to anthropogenic ecological and demographic changes (*36, 39, 41, 42*) including mosquito-friendly habitats via open-pit mining in the Guyana shield (*2*) may not yet be detectable in population genetic data.

We detected thirteen large (3-19 Mb) segregating haplotype blocks — likely chromosomal inversions (*43*) — showing perfect linkage disequilibrium among physically distant polymorphisms (fig. S8, fig. S9; table S3). The two haplotype lineages (presumed opposite orientations) of an inversion typically show elevated sequence divergence from each other as well as substantial diversity within themselves (fig. S10). There are few examples of alternate inversion haplotypes being fixed across populations, except in lower-diversity populations like Pará in which genetic drift presumably plays a larger role (fig. S9). Heterozygosity varies considerably both among samples and across the genome (Fig. 3C), including copy number variants overlapping 6425 genes (table S4), which also vary by population (fig. S11).

## Positive selection signals may reflect emerging insecticide resistance

Numerous signals of positive selection are apparent across these populations (tables S5, S6, and S7). Genome-wide F_ST_ scans between adjacent populations reveal the same 150 kb window (2RL_8.425-8.575Mb) within Inversion B to be an outlier, sometimes the top outlier, in multiple independent comparisons (Fig. 4A). G1 and G123 statistics for several populations are also outliers in this window (Fig. 4B), which overlaps 20 genes including a cluster of six cytochrome P450 genes (Fig. 4C). Even a single incidence of this selection signal would be striking, as in Peru (Fig. 4B) or Guainía (fig. S12A), yet this window is among the top outliers in approximately half our populations by at least one metric (Fig. 4D). In some populations, the selection signal is clearly concentrated on the P450 genes, especially *CYP6AA1* (*ADAR2_008159*), tied to pyrethroid resistance in *A. funestus* (*44*) and *A. gambiae* (*45*). The top F_ST_ value between Peru and Amazonas is in *CYP6AA1*, and missense variant *CYP6AA1* T283K is one of the two highest F_ST_ values in this window between Cacao and other French Guiana populations. In Guainía, similarly high F_ST_ values occur across a wider window (2RL_8-9Mb) and many individuals exhibit near-zero heterozygosity across several megabases surrounding this genomic region (Fig. 3C; fig. S12A), although G1 peaks within the narrower window at 2RL_8.465Mb. Various derived alleles within Inversion B have been selected, rather than the entire inversion haplotype, and these alleles are not closely related across populations (fig. S12B), suggesting soft sweeps on distinct genetic backgrounds rather than a single species-wide hard sweep of a novel adaptive allele.

**Fig. 4.**
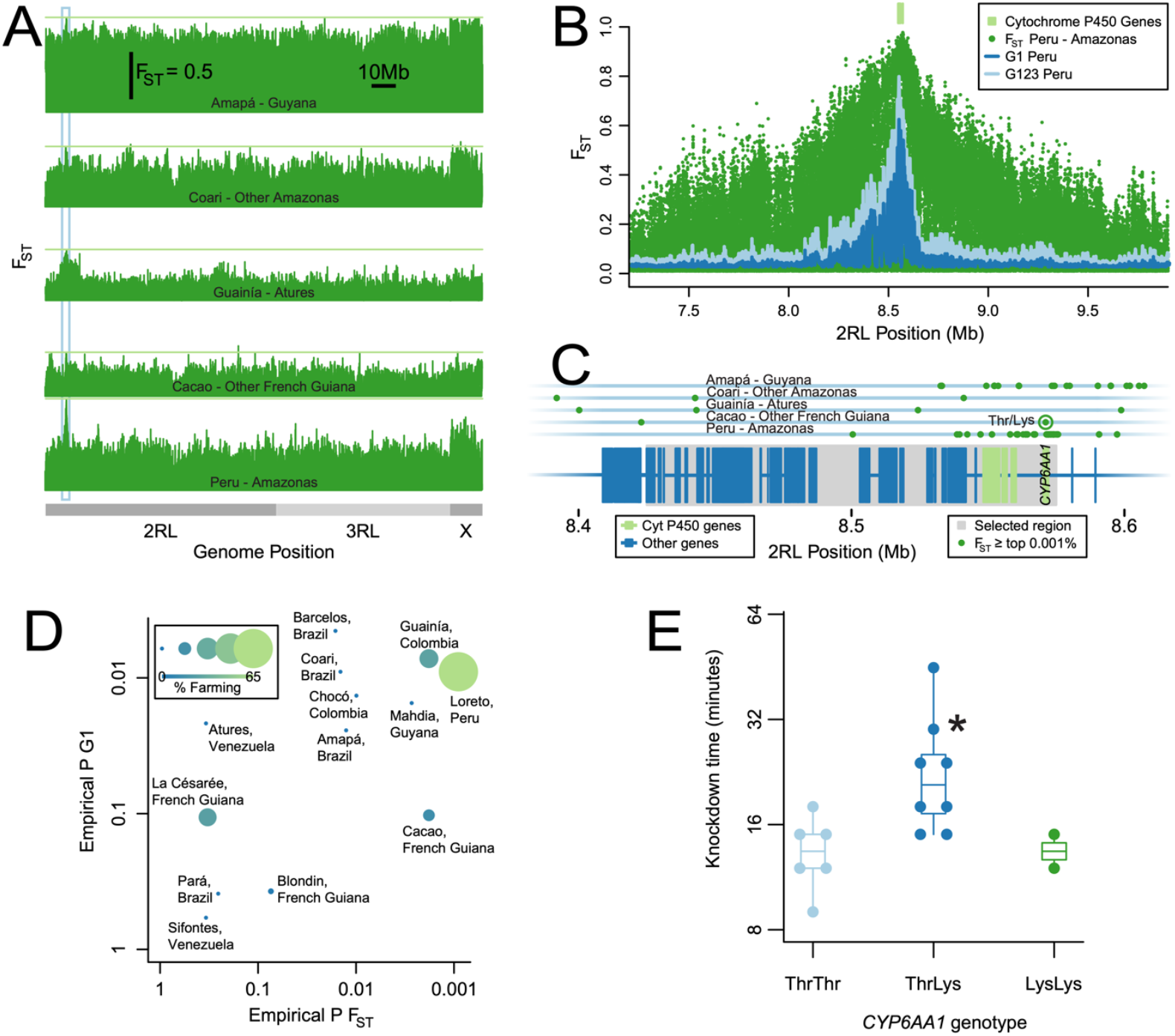
Signals of selection in a P450-dense genomic region. **(A)** In genome-wide F_ST_ scans of five independent pairwise comparisons, the same region (blue-outlined box) has the highest or nearly-highest F_ST_ (light green horizontal lines). **(B)** Peru-Amazonas F_ST_ (dark green), Peru G1 (dark blue), and Peru G123 (light blue) all peak around 2RL 8.5 Mb, near a cluster of cytochrome P450 genes (light green). **(C)** Genes in the 150 kb region (2RL_8.425-8.575Mb, grey box) showing a positive selection signal in multiple populations. Top F_ST_ outliers (0.001% threshold) from the five comparisons (A) are plotted (dark green dots). The *CYP6AA1* Thr283Lys polymorphism is circled. **(D)** For each population, the top values of G1 and F_ST_ (against genetically similar sister population(s)) in the 2RL_8.425-8.575 Mb window were ranked among all autosomal 150 kb windows, yielding empirical P values. Positive selection is indicated by low P (meaning most windows have lower G1 or F_ST_). Size and color of points indicate the proportion of surrounding land devoted to “mosaic of farming uses”, which is high for several strongly-selected populations. **(E)** Knockdown bottle assay with 12.5 µg/mL deltamethrin of wild-caught Cacao *A. darlingi* genotyped at *CYP6AA1* Thr283Lys polymorphism. Heterozygotes survive significantly longer (P < 0.05).

Given the well-established link between cytochrome P450 genes and toxin resistance (*23, 44*), we assessed whether the observed selection signals could be tied to land use practices near our collection sites that might expose mosquitoes to insecticides. Runoff of agricultural pesticides into Latin American aquifers (*46*) could be the main source of exposure, as hypothesized for resistance in African *Anopheles* (*47*) and possibly other non-targets of pesticides (*48*). We ranked the 2RL_8.425-8.575Mb window against all autosomal 150 kb windows based on F_ST_ and G1 to generate empirical P values (Fig. 4D). The lowest empirical P values (< 0.002) occurred for F_ST_ in Peru, Cacao, and Guainía, and two of these (Peru and Guainía) also had among the lowest empirical P values for G1 (< 0.01). Notably, all three of these locations, unlike most of our collection sites, have substantial land use dedicated to “mosaic of farming uses” (Fig. 1B), although there is no significant correlation between proportion of farming land use and selection signals (P > 0.1).

To test if selected alleles impact insecticide resistance, we collected sixteen wild *A. darlingi* from Cacao, French Guiana and tested their resistance in a bottle assay with 12.5 µg/mL deltamethrin (Fig. 4E). We genotyped these mosquitoes at *CYP6AA1* T283K and found knockdown time of TK heterozygotes (median = 21 minutes) was significantly longer than that of either ancestral TT (median = 13.5 minutes) or derived KK (median = 13.5 minutes) homozygotes (P < 0.05). There was no significant phenotypic difference between homozygotes (P > 0.1). We note that our sample size was small with only two KK homozygotes, the overdominant genotypic effect on resistance is not straightforward to interpret, our results merely identify T283K as linked to the phenotype rather than causal, and mosquitoes of all genotypes were killed within an hour.

Other regions of the genome show selection signals beyond this window, though not as strong or as consistently across populations (figs. S13, S14, and S15; tables S5, S6, and S7). F_ST_ for Cacao versus other French Guiana was highest in a different cluster of P450 genes at 2RL_79.3 Mb (fig. S15A), peaking at *CYP6M2* (*ADAR2_008623*) tied to pyrethroid-resistance in *A. gambiae* (*49*). This double P450 signal strengthens our conclusion that the P450s themselves, and not unrelated linked genes, are the targets of selection. A window at 2RL_93.635-93.785 Mb, containing a set of seven ATP-binding cassette sub-family G genes, is a top F_ST_ outlier in two independent pairwise comparisons: between French Guiana sites and between Amazonas, Brazil sites (fig. S13; table S5). A strong selection signal centered on 2RL_49.950-50.050 Mb is implicated by F_ST_, G1, and G123 in Barcelos (fig. S14). The top genome-wide F_ST_ for this comparison occurs in *Keap1* (*ADAR2_006983*), which also shows signals of positive selection in *A. gambiae* (*50*) and *A. arabiensis* (*51*), plausibly because of its role in a metabolic insecticide resistance pathway (*22*).

We do not observe notable positive selection at the orthologs of target-site resistance genes known in other *Anopheles*. In fact, target codons L995 in *vgsc* (site of the *kdr* mutation), A296 in *rdl*, and G280 in *ace1* (*24*) exhibit no nonsynonymous variation (fig. S15B), consistent with previous results (*25*). P450s and *Keap1* underlie metabolic resistance, suggesting that this mechanism has been favored in *A. darlingi* over target-site mechanisms of resistance. While the specific adaptive polymorphisms remain unidentified, it is notable that there is very little evidence that gene duplications of P450 or *Keap1* are selection targets, as hypothesized for *A. gambiae* (*45*) and *A. funestus* (*52*). Copy number variants in *CYP6AA1, CYP6M2*, and adjacent P450 genes (table S4; fig. S11A) are rare and not associated with the selected alleles. Some populations with selection signals had no or very few P450 duplications, and these were uncommon in samples homozygous for the derived allele at high-F_ST_ variants. We observed no *Keap1* duplicates.

Our results have several implications for vector control across Latin America. *A. darlingi* is extremely diverse and likely harbors latent potential to adapt to almost any novel intervention designed to kill it. Signals of positive selection suggest that this may already be occurring and indicate that broad continuing assays for emerging insecticide resistance across the continent are warranted. Although long-distance gene flow is limited, that may not matter as resistance can evolve in parallel on different genetic backgrounds across habitats, as has seemingly occurred with P450s. Continued population genomic monitoring can inform both disease control efforts and broad conclusions about ecological and evolutionary diversity in neotropical vectors.

## Acknowledgments

We thank Nora Besansky, Romuald Carinci, Auden Cote-L’Heureux, Carlo Maria De Marco, Tatiane M. P. de Oliveira, Silvia Díaz, Elder Figueira, Pascal Gaborit, Zachary Johnson, Cheyenne Knox, Mara Lawniczak, Tovi Lehmann, Martin Llewellyn, Aleksei Makunin, Nelson Moncada, Caio C. Moreira, Monique A. Mota, Maycon Neves, Rachel Newmiller, Ruchit Panchal, Seth Rudman, Philipp Schwabl, Meg Shieh, Nithya Swaminathan, Stanislas Talaga, Ana Villarroel, and malaria control staff and local residents at various collection locations.

## Author contributions

Conceptualization: JEC, DEN

Data curation: JAT, RB, AME

Formal analysis: JAT, NJA, ERL, MCC

Funding acquisition: MAMS, MG, JEC, DEN

Investigation: KAK, MAMS, JEC Methodology: JAT, DEN

Project administration: ML, DEN

Software: SN

Resources: EC, SB, DG, ESB, JEM, MS, RN-R, MAMS, MEG, MLQ, HC, MG

Supervision: AME, DEN Validation: J-BD

Visualization: JAT

Writing – original draft: JAT

Writing – review & editing: JAT, MEG, MAMS, JEC, DEN

## Data and materials availability

All raw sequencing data has been deposited at NCBI SRA, BioProject PRJNA1169887. Genotypes are accessible at Malaria Genome Vector Observatory. Transfer of samples was made possible by a material transfer agreement between Universidade de São Paulo and Harvard: MTA No 1017829, SISGEN Registration no. RCD3CF5, Export permit IBAMA no. 24BR0490030/DF.

## Supplementary Materials

Materials and Methods

Figs. S1 to S15

Tables S1 to S7

References (*53*–*70*)

