## Supplementary Methods and Figures for "Population genomics of *Anopheles darlingi*, the principal South American malaria vector mosquito"

**The PDF file includes:**

Materials and Methods
Figs. S1 to S15

**Other Supplementary Materials for this manuscript include the following:**

Tables S1 to S7

### Materials and Methods

We collected 1094 adult female specimens of *A. darlingi* from 16 locations in Colombia, Venezuela, Peru, Brazil, Guyana, and French Guiana (table S1) for whole-genome-sequencing, drawing from new and published (e.g. Chabanol et al. 2024) collections. We extracted DNA from whole mosquito carcasses using either the Qiagen DNeasy Blood & Tissue Kit or the MagBio HighPrep Blood & Tissue DNA Kit with Buffer C (Makunin et al. 2022). For some samples, we performed automatic DNA extraction with the KingFisher Duo Prime system (Thermo Scientific). We used Nextera low-input sequencing libraries to sequence whole genomic DNA with 151 bp paired-end reads on an Illumina NovaSeq instrument at the Broad Institute. All raw sequencing data has been deposited at NCBI SRA, BioProject PRJNA1169887.

Additionally, in May and July 2022, with four Mosquito Magnet® traps baited with octenol, we collected sixteen wild adult female *A. darlingi* from Cacao, French Guiana for individual insecticide phenotyping. We subjected individual insects to a bottle assay with 12.5 µg/mL deltamethrin and measured knock-down time over one hour. We used t-tests to assess phenotypic differences among genotypes.

We confirmed species identity by aligning reads to *COX1* sequences from *A. darlingi* and other mosquito species, and by searching reads for the species-specific *ITS2* kmer CTGCGTACTGATGATTTGATTGACGCGCCGCGTCCCCAA. All reads were aligned to the *A. darlingi* idAnoDarlMG_H_01 reference genome (NCBI Accession: GCF_943734745.1) using bwa-mem2 v.2.2.1 (Vasimuddin et al. 2019; command: bwa-mem2 mem -M), Picard MarkDuplicates v.2.26.10 and samtools v.1.16.1 (Danecek et al. 2021; commands: samtools view, samtools sort, samtools index).

Single nucleotide polymorphisms (SNPs) and indel variants were called using GATK HaplotypeCaller v.4.3.0.0 (McKenna et al., 2010; Poplin et al. 2018) using SNP and indel heterozygosity parameters from the Anopheles gambiae 1000 Genomes project (Anopheles gambiae 1000 Genomes Consortium 2017). SNPs were hard-filtered with QD < 2, FS > 60, ReadPosRankSum < -8, QUAL < 30, SOR > 3, MQ < 40, and/or MQRankSum < -12.5. Indels were hard-filtered with QD < 2, QUAL < 30, FS > 200, and/or ReadPosRankSum > -20.

We examined 49,816,420 total segregating sites. We identified potentially contaminated samples by examining the relative depth of each allele within heterozygous genotypes: while this ratio is close to 1:1 for most sites in most single individuals, a peak at a lower minor allele depth reveals the putative contaminants, which we confirmed to have unusually high heterozygosity compared to other samples from the same location and subsequently ignored (fig. S1). Allele frequencies (e.g. for $F_{ST}$) were calculated from all remaining samples. For analyses that required accurate genotypes (e.g. PCA and estimates of heterozygosity), we restricted analysis to samples with median depth ≥8x (N = 815, table S1) and counted genotypes with <5x coverage to be missing. We performed statistical tests and data visualization in R (v. 4.3.3; R Core Team, 2024). We conducted principal component analysis (PCA) with the princomp function in R for various subsets of the data, restricting analysis to sites with no missing data across all included samples. We performed ADMIXTURE analysis (Alexander et al. 2009) on a set of 154,930 variants spaced at least 1 kb apart and excluding variants within inversions, with K ranging from 1 to 10, choosing the value of K with the lowest cross-validation error. We conducted interpopulation phylogenetic analysis with TreeMix (Pickrell and Pritchard 2012), restricting migration edges at zero. Genomic analysis was facilitated by VectorBase (Giraldo-Calderón et al., 2015). We used RAxML with -m GTRCAT for phylogenetic analysis of haplotypes from inversions and selection regions (Stamatakis 2014).

We calculated population genetic statistics π and Tajima's D with Perl scripts incorporating Perlymorphism (Bio::PopGen, BioPerl version 1.007000; Stajich & Hahn, 2005). For calculation of $F_{ST}$ we used FstPerSite.pl and other scripts at https://github.com/jacobtennessen/MalariaHallmarks/ (Tennessen and Duraisingh 2021). For assessing $F_{ST}$ at individuals sites (e.g. for outlier selection signals) we used Weir and Cockerham's estimator (Weir and Cockerham 1984), while for mean pairwise differences we replicated the methods used for the *Anopheles gambiae* 1000 Genomes project (Anopheles gambiae 1000 Genomes Consortium 2017) based on Bhatia et al. (2013): the ratio of averages for Hudson's $F_{ST}$ estimator, only considering variants segregating in both populations and excluding inversions. For comparison with *A. gambiae* and *A. coluzzii*, we obtained published $F_{ST}$ estimates between continental populations (Anopheles gambiae 1000 Genomes Consortium 2020), excluding islands Bioko and Mayotte which could have distinct bottleneck histories as well as the "uncertain" populations like Kenya that could not be assigned to one of these two species. We considered it a signal of selection-driven elevated $F_{ST}$ when we observed at least three autosomal variants with $F_{ST}$ greater than ten times the mean autosomal $F_{ST}$ for that comparison, spanning at least 500 bp (for representation on separate read pairs), but with no more than 200 kb between adjacent variants (though the full region under selection may extend over 200 kb). We calculated G1 and G123 (Harris et al. 2018) on autosomes using code written for this purpose (https://github.com/sanjaynagi/ad1000g-selection/blob/main/darlingi-g123.ipynb), with window size of 400 and step size of 200, and considered the top 50 G1 or G123 windows in each population to be outliers. Thus, for both $F_{ST}$ and G1/G123, we only considered selection signals on autosomes, since they could be compared to a large number of putatively neutral background sites. The X in contrast may have a different effective size and evolutionary rate precluding direct comparison with the autosomes, and because it is small and a single inversion (M, table S3) extends for more than half its length, it is more challenging to distinguish adaptive signal from neutral variation.

We identified inversions as sets of at least 25 polymorphisms spanning at least 2 Mb with minor allele frequency at least 5% in one population and all showing perfect linkage disequilibrium with each other, either across all populations or within one of the major clades (Chocó, Orinoco, or Amazon/Atlantic). Boundaries of inversions were defined by calculating $F_{ST}$ between alternate homozygotes from the same population, which tended to reveal sharp edges (fig. S8) that closely matched the range of the variants in perfect linkage disequilibrium. Lacking a direct way to match our observed inversions with those reported from polytene chromosomes (Cornel et al. 2016), we did not follow the same naming scheme and assigned a capital letter to each inversion.

Copy number variants were discovered following the pipeline used for *A. gambiae* (Lucas et al. 2019). Coverage was calculated in non-overlapping 300bp windows, and each window was normalized by the median coverage value for all autosomal windows with the same GC content. After removing windows in which >50% of reads were aligned with mapping quality 0, and windows for which the percentage GC content was rarely represented within the genome (fewer than 100 bins), the copy number state at each window in each individual was then estimated using a Gaussian HMM applied to the normalized coverage, implemented in Python's hmmlearn package, with copy number states ranging in unit increments from 0 to 12, transmission probability of $10^{-5}$, and emission variance per copy determined for each sample as the normalized autosomal coverage variance. The copy number of each gene was calculated as the mode of the copy number state across the windows covering that gene. Given the relatively high coverage variance observed among low-coverage samples, we only examined copy number variants among samples with median depth ≥ 15x.

We used MapBiomas (Souza et al. 2020) to identify land use categories within 5 km of sample collection sites. We used the "Mosaic of Farming Uses" category as a proxy for potential insecticide exposure. We tested for a correlation between "Mosaic of Farming Uses" and the minimum of the empirical P values of $F_{ST}$ and G1 for the 2RL_8.425-8.575Mb window at each collection site, using a Wilcoxon signed-rank test.

To identify bloodmeal DNA, we aligned reads to complete mitochondrial genomes of several vertebrates, including human, dog, cow, pig, horse, chicken, and duck. We discovered that mosquito reads did align to some portions of these genomes, so we masked those regions and for subsets of reads aligning elsewhere, we used BLAST and manual alignment to confirm the vertebrate identity of the reads. To test if human DNA could derive from handling, our French Guiana mosquitoes were evaluated visually for abdominal blood and we confirmed that detection of human DNA was positively correlated with this visual assessment (Fisher's exact text, $P < 0.05$), suggesting it truly reflects blood meals. For Plasmodium DNA, we employed a similar approach by aligning reads to the complete genomes of *P. falciparum* and *P. vivax*, before confirming with BLAST and manual alignment. We tested whether *Plasmodium* spp. DNA was significantly more common within samples positive for human DNA using Fisher's exact text.

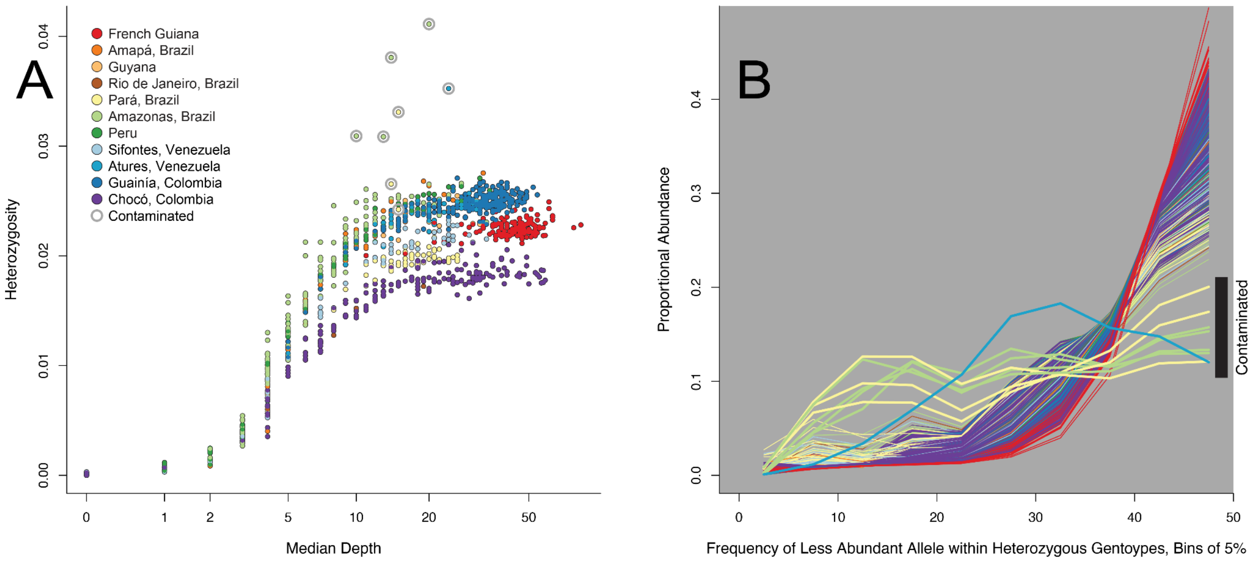


**Fig. S1**.

**Heterozygosity tests for contaminated samples. (A)** Heterozygosity plotted against median depth for all samples. Heterozygosity increases with depth before plateauing around 10x coverage and varies among populations. Contaminated samples (circled in grey) are apparent as showing unusually high heterozygosity given their depth and population. (**B)** Histogram for each sample of the relative depth of the less abundant allele within heterozygous sites (colored by population as in A). "Less abundant allele" refers to frequency among reads within that one sample, not the population-wide minor allele. For most samples, most heterozygous sites have both alleles close to 50% as expected, but the eight contaminated samples (thicker lines) have a left-shifted distribution suggesting a large proportion of sites with skewed allele depths.

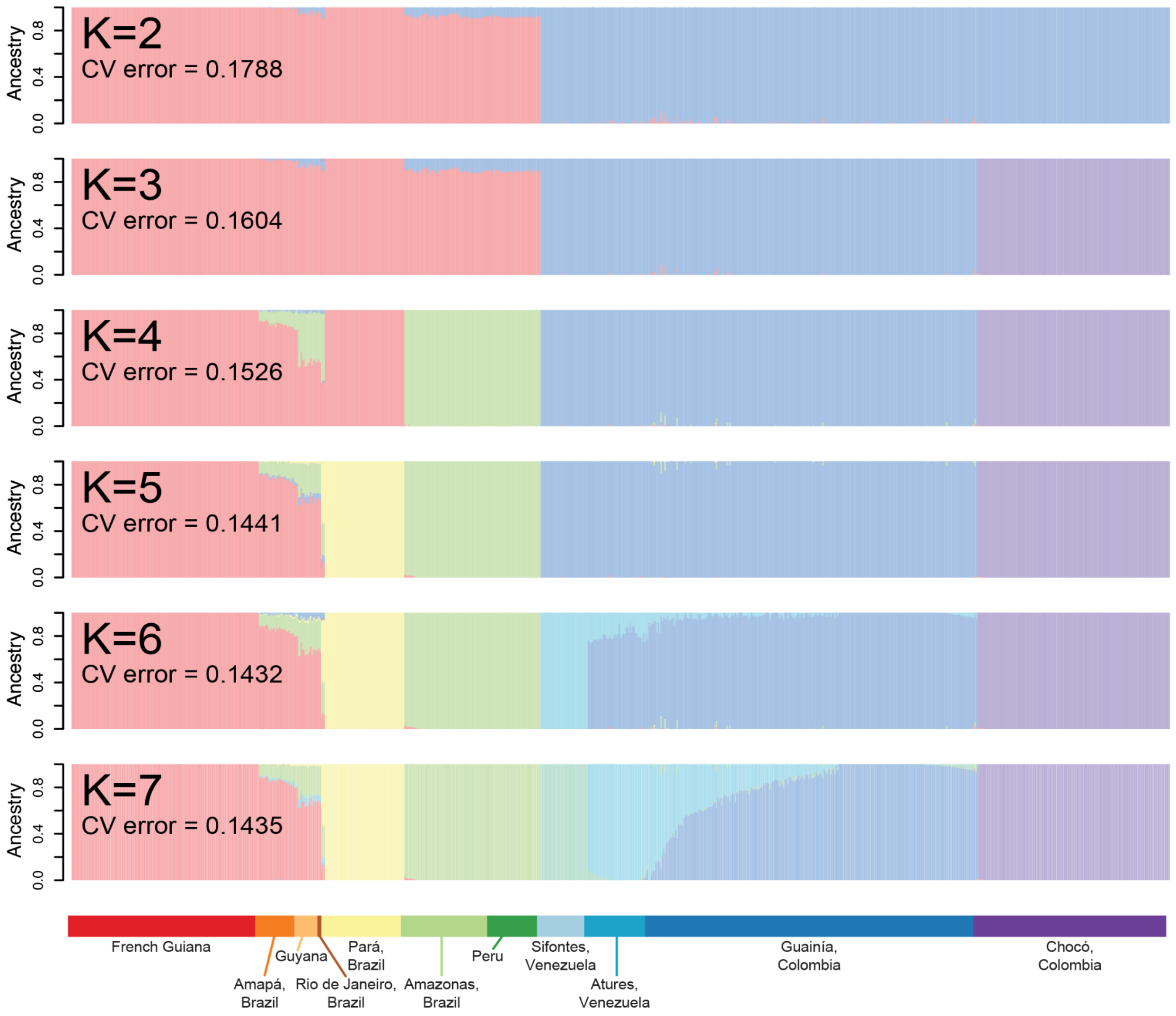


**Fig. S2.**

**Ancestry assignment via ADMIXTURE for values of K less than the optimal value of 8**. The fundamental division (K=2) is between Amazon/Atlantic populations and Orinoco/Pacific populations, followed by a split into the three major clades (Amazon/Atlantic, and Orinoco, and Pacific) at K=3. Cross-validation (CV) error declines with increasing K until K=8 (Fig. 2A) with a CV error of 0.1422, after which CV increases.

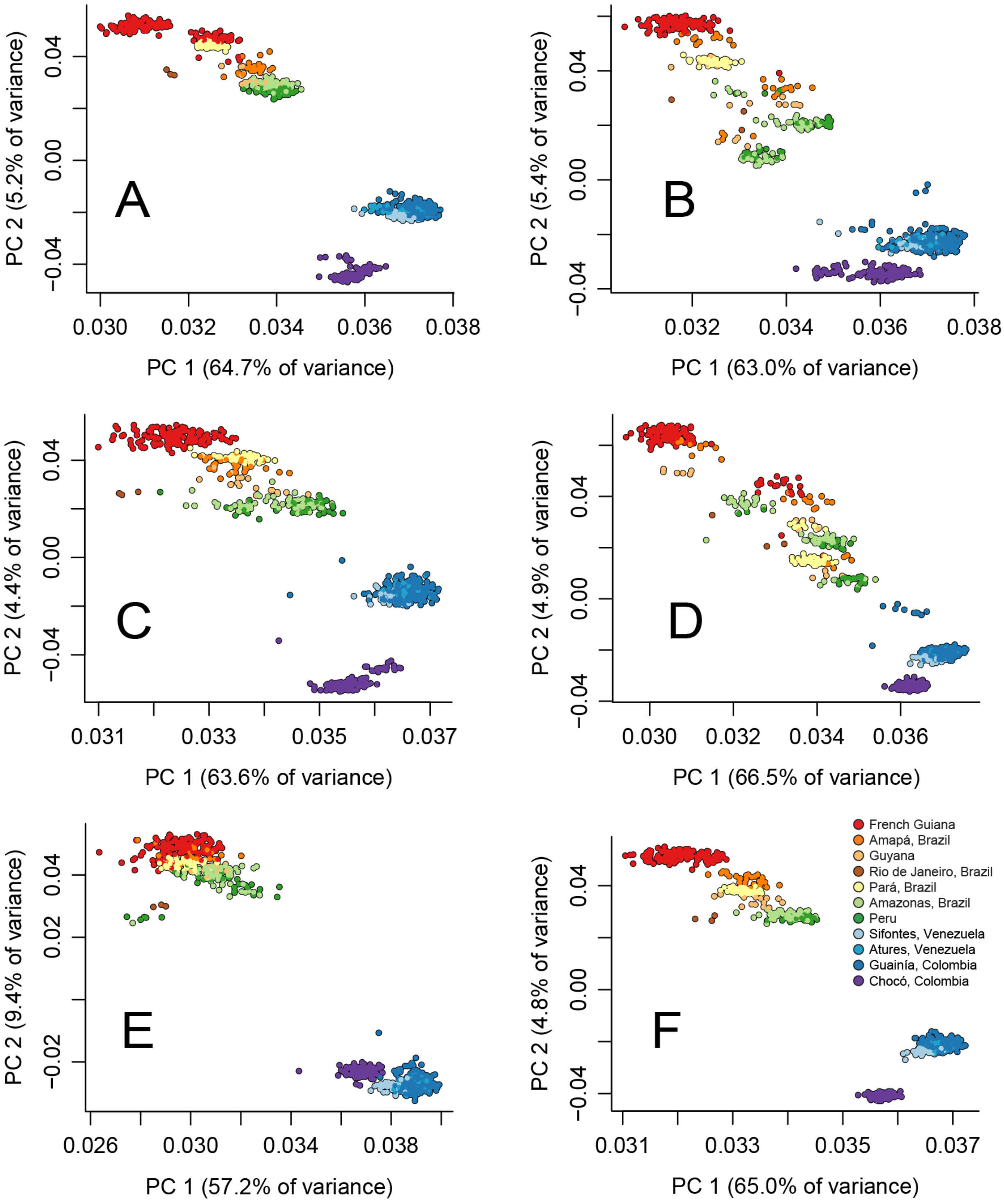


**Fig. S3.**

**Principal component analysis using subsets of the genome.** Colored as in Fig. 2B. (**A**) Chromosome 2RL, upstream of centromere. (**B**) Chromosome 2RL, downstream of centromere. (**C**) Chromosome 3RL, upstream of centromere. (**D**) Chromosome 3RL, downstream of centromere. (**E**) Chromosome X. (**F**) Full genome excluding all inversions.

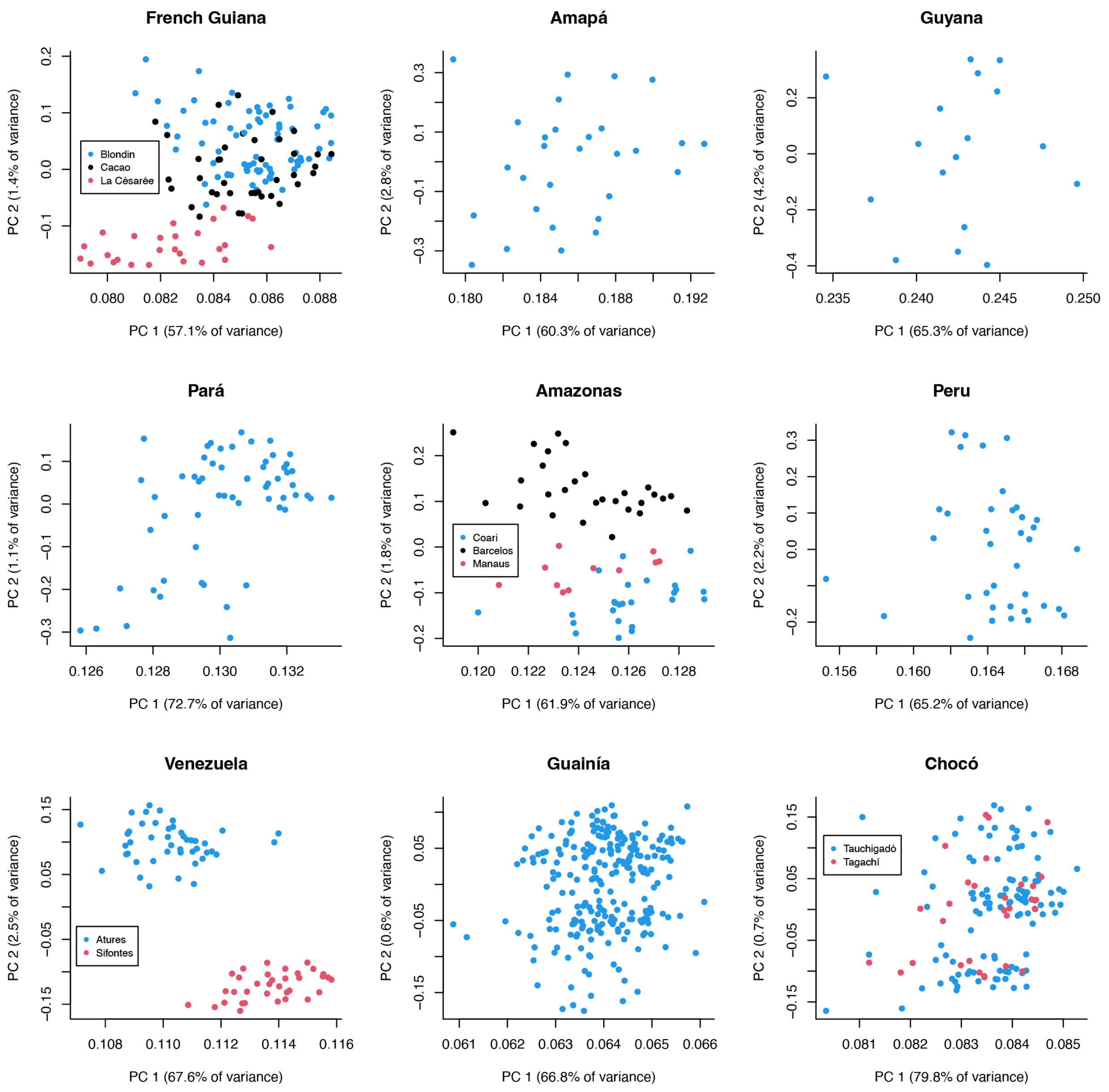


**Fig. S4.**
**Principal component analysis for distinct geographical locations**. When more than one distinct collection site is included together, locality is indicated by color.

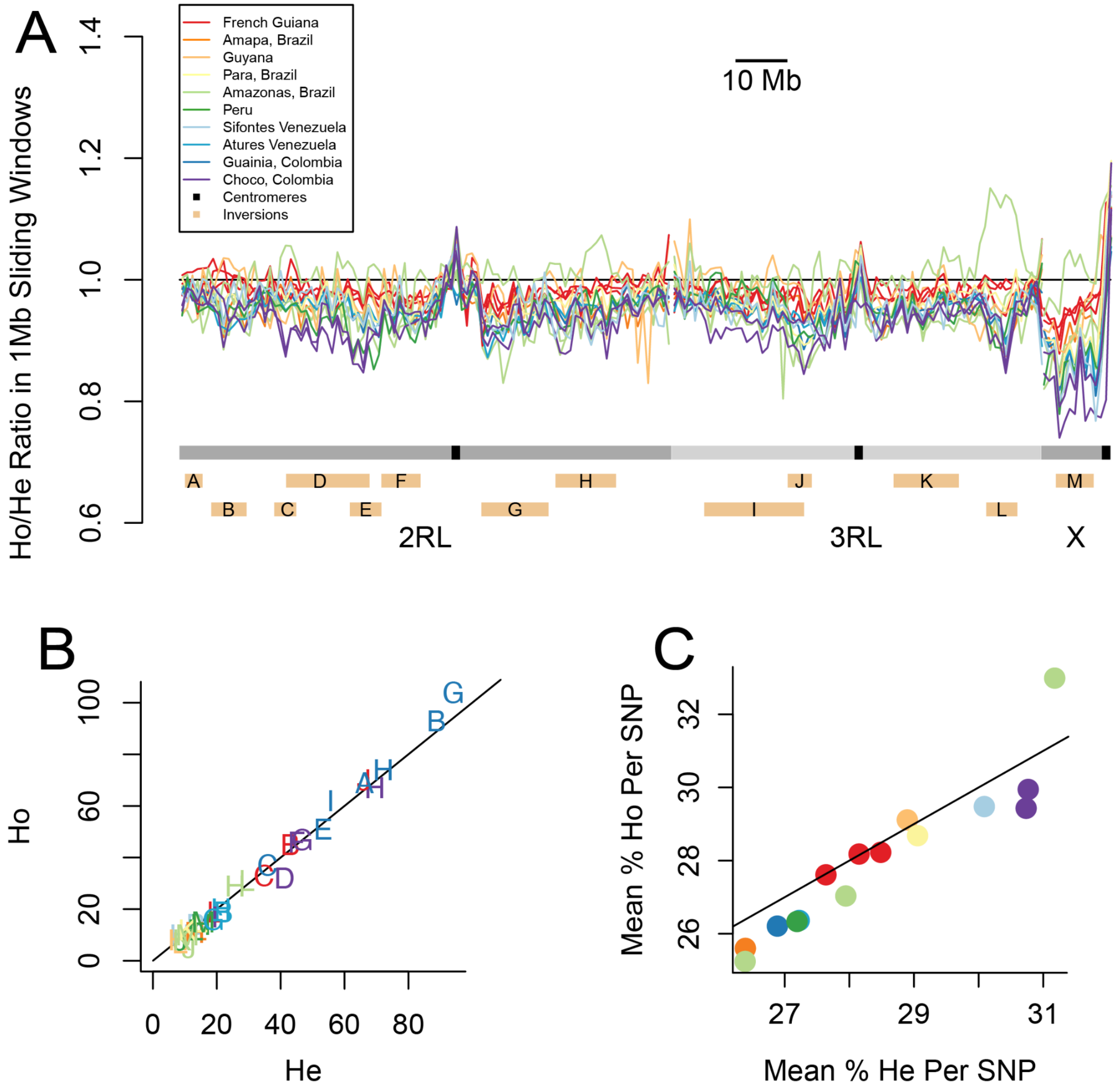


**Fig. S5.**

**Assessment of Hardy-Weinberg equilibrium across the genome. (A)** Logarithmic mean of $(H_o + 1)/(H_e + 1)$ in 1 Mb windows for polymorphisms with population-specific minor allele frequency ≥ 5%, where $H_o$ = count of observed heterozygotes per population and $H_e$ = count of expected heterozygotes per population. Most samples from Manaus, Amazonas are heterozygous for inversion L, causing a minor peak, but the excess is not significant with this small sample size. **(B)** $H_o$ versus $H_e$ for all inversions within populations where they are common. Inversions are labeled with capital letters and populations are indicated by color, as in (A). **(C)** Mean percentage of observed homozygous versus expected homozygous individuals in each population, for single nucleotide polymorphisms (SNPs) with population-specific minor allele frequency ≥ 5%. Colors indicate population as in (A). Note that these values are per SNP (a different number for each population), not per site, so they do not reflect nucleotide diversity (π).

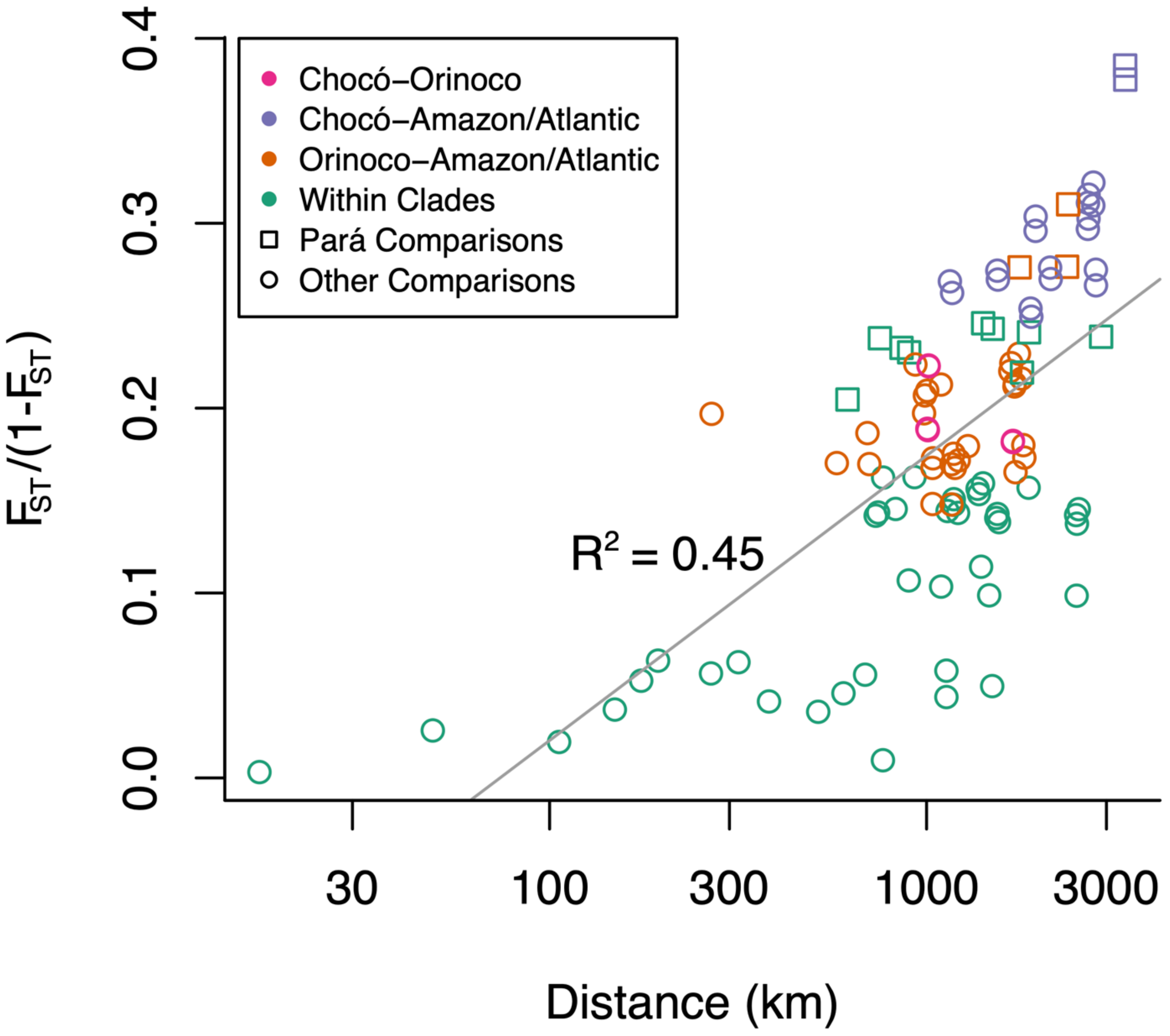


**Fig. S6**.

**Pairwise genetic distance as a function of geographic distance.** Pairwise values of $F_{ST}/(1- F_{ST})$ among fifteen *A. darlingi* collection sites (Rio de Janeiro excluded for sample size) are shown, colored by clade composition (Fig. 2C). Genetic distance is correlated with geographic distance ($R^2 = 0.45$), though most of this effect can be attributed to between-clade $F_{ST}$ exceeding within-clade $F_{ST}$. Comparisons involving Pará are indicated with squares; $F_{ST}$ against Pará tends to be larger than other comparisons, presumably because Pará has experienced unusually high genetic drift, explaining its relatively low diversity (Fig. 3A).

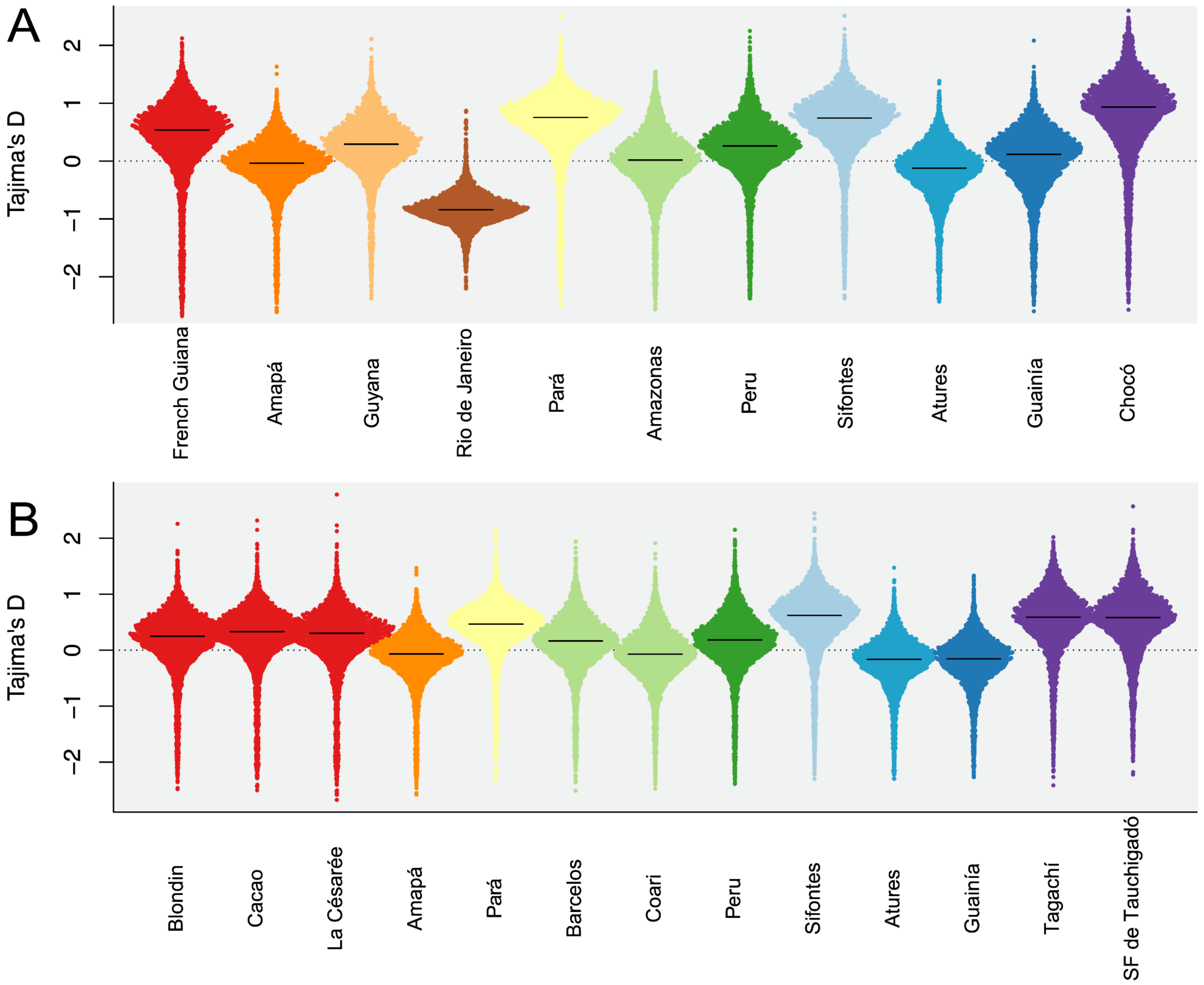


**Fig. S7.**
**Tajima's D in 20 kb windows per collection locality.** Medians indicated by horizontal lines. **(A)** Full populations, all individuals with depth ≥8x. **(B)** Subsamples of 25 mosquitoes from each of the specific collection sites, for collection sites with at least 25 individuals with depth ≥8x.

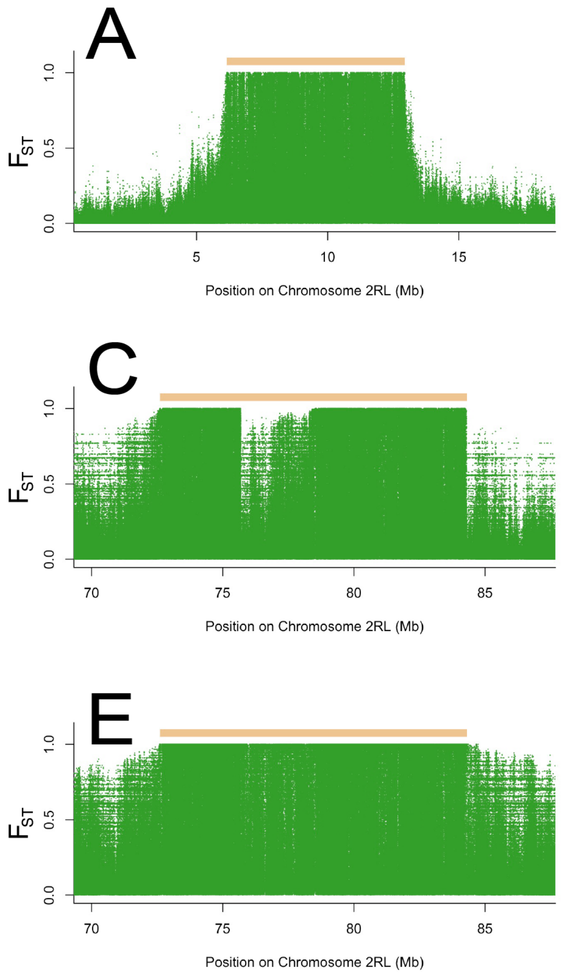


**Fig. S8**.

**Boundaries of inversions revealed as "tepuis" of $F_{ST}$ between homozygotes for alternate haplotypes.** Inversion boundaries are consistent among clades (Amazon/Atlantic, Orinoco, and Pacific). Green dots indicate $F_{ST}$ values per variant, and tan rectangles indicate inferred inversion positions. **(A)** $F_{ST}$ between homozygotes for Inversion B in Guainía and Venezuela. **(B)** $F_{ST}$ between homozygotes for Inversion B in French Guiana, Amapá, and Guyana. **(C)** $F_{ST}$ between homozygotes for Inversion H in Guainía and Venezuela. Note the region of low $F_{ST}$ within the inversion, possibly indicating a translocation. **(D)** $F_{ST}$ between homozygotes for Inversion H in Chocó. Note the same putative translocation observed in (C). **(E)** $F_{ST}$ between homozygotes for Inversion H in French Guiana, Amapá, and Guyana. **(F)** $F_{ST}$ between homozygotes for Inversion L in Peru and Amazonas.

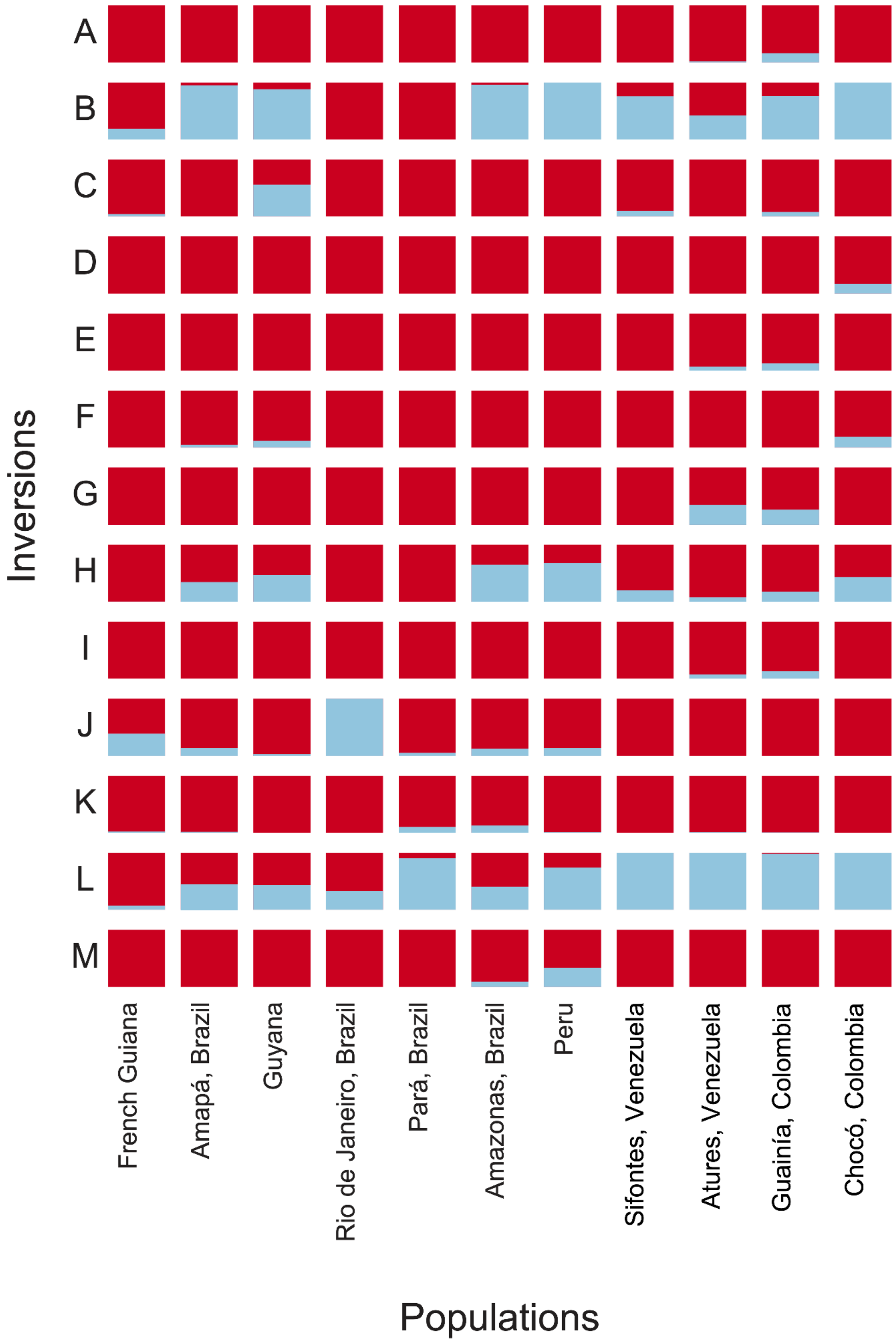


**Fig. S9**.
**Frequencies of inversion haplotypes across populations.** Arbitrarily colored blue or red.

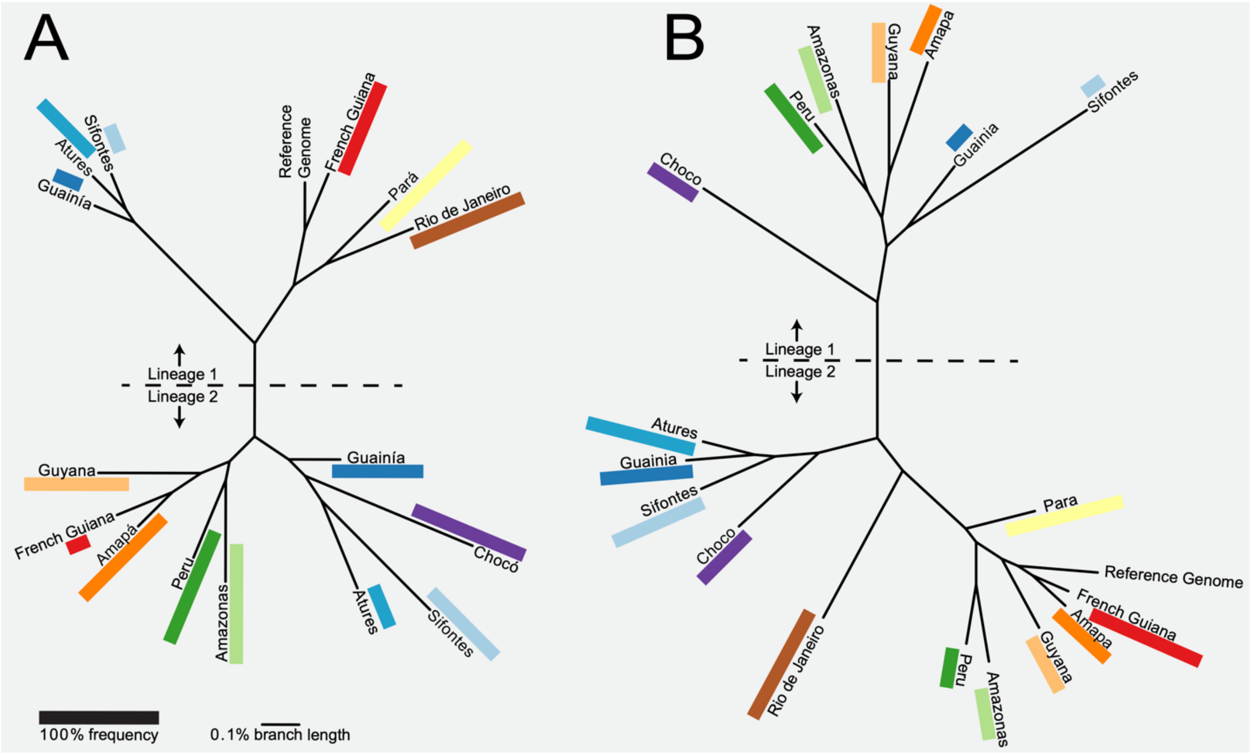


**Fig. S10**.

**Phylogeny of representative homozygotes for inversions.** Lengths of solid rectangles are proportional to the frequency of each haplotype in each population, and a dotted line separates the two main haplotype lineages (presumed opposite orientations). **(A)** Inversion B. **(B)** Inversion H.

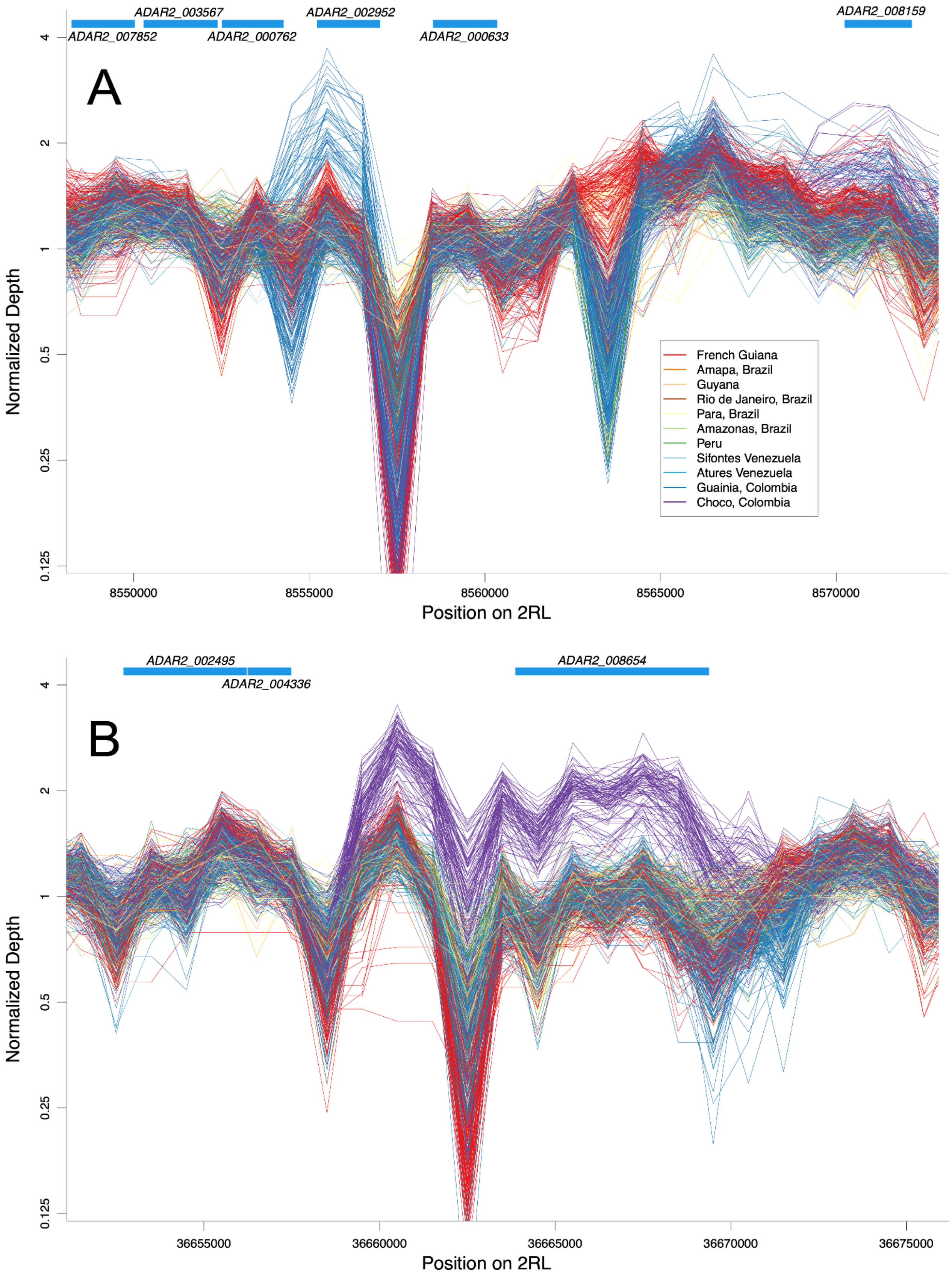

A
ADAR2_003567
ADAR2_002952
ADAR2_008159
ADAR2_007852
ADAR2_000762
ADAR2_000633
Normalized Depth
4
2
1
0.5
0.25
0.125
French Guiana
Amapa, Brazil
Guyana
Rio de Janeiro, Brazil
Para, Brazil
Amazonas, Brazil
Peru
Sifontes Venezuela
Atures Venezuela
Guainia, Colombia
Choco, Colombia
8550000
8555000
8560000
8565000
8570000
Position on 2RL
B
ADAR2_002495
ADAR2_004336
ADAR2_008654
Normalized Depth
4
2
1
0.5
0.25
0.125
36655000
36660000
36665000
36670000
36675000
Position on 2RL

**Fig. S11.**
**Examples of copy number variants.**
Each line represents a sample with median depth ≥ 15x and shows coverage in 1 kb windows, normalized to the mean coverage for that sample. Samples are colored based on geography. Duplications are visually evident in the higher depth of certain samples in certain windows. Genes are shown as blue rectangles at the top of each plot. **(A)** Cluster of P450 genes at 2RL_8.5 Mb. A duplication predicted by our HMM (table S4) in 15% of samples from Guainía, Colombia (dark blue) overlaps gene *CYP6P5* (*ADAR2_002952*), while another duplication in 39% of samples from Chocó, Colombia (purple) overlaps gene *CYP6AA1* (*ADAR2_008159*). **(B)** A duplication at 2RL_36.66 Mb predicted in a majority of samples from Chocó, Colombia (purple) but not in any samples from other locations, overlapping gene *ADAR2_008654* (uncharacterized).

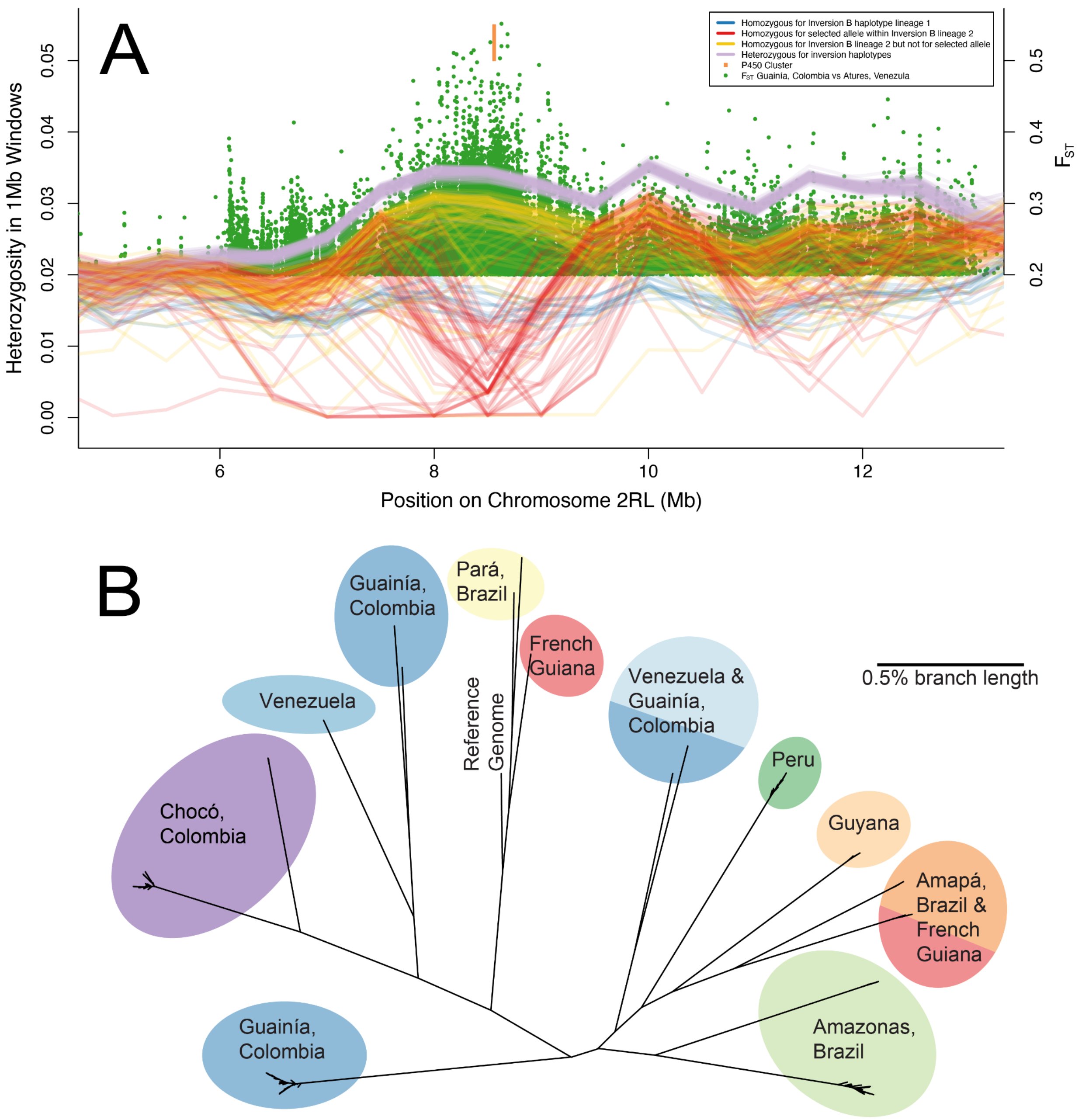


**Fig. S12.**

**Additional details of selection signal at 2RL_8.425-8.575Mb (Fig. 4). (A)** Selection signal in Guainía, Colombia, with heterozygosity per sample in sliding 1Mb windows (lines, from Fig. 3C) and $F_{ST}$ versus Atures, Venezuela (green dots, from Fig. 4A). Samples are colored based on haplotype at Inversion B and genotype at position 2RL:8434067 where the non-reference allele is a proxy for the selected haplotype. Homozygotes for the selected allele at 8434067 (red) tend to show a wide swath of reduced heterozygosity. **(B)** Phylogeny of the 100kb window 2RL:8470-8570kb, for the 222 samples that have at least 8x genome-wide depth and heterozygosity <0.001 in this region and are thus putatively homozygous for a selected allele. Substantial divergence among populations suggests that unrelated alleles on different genetic backgrounds have been selected in different geographic regions.

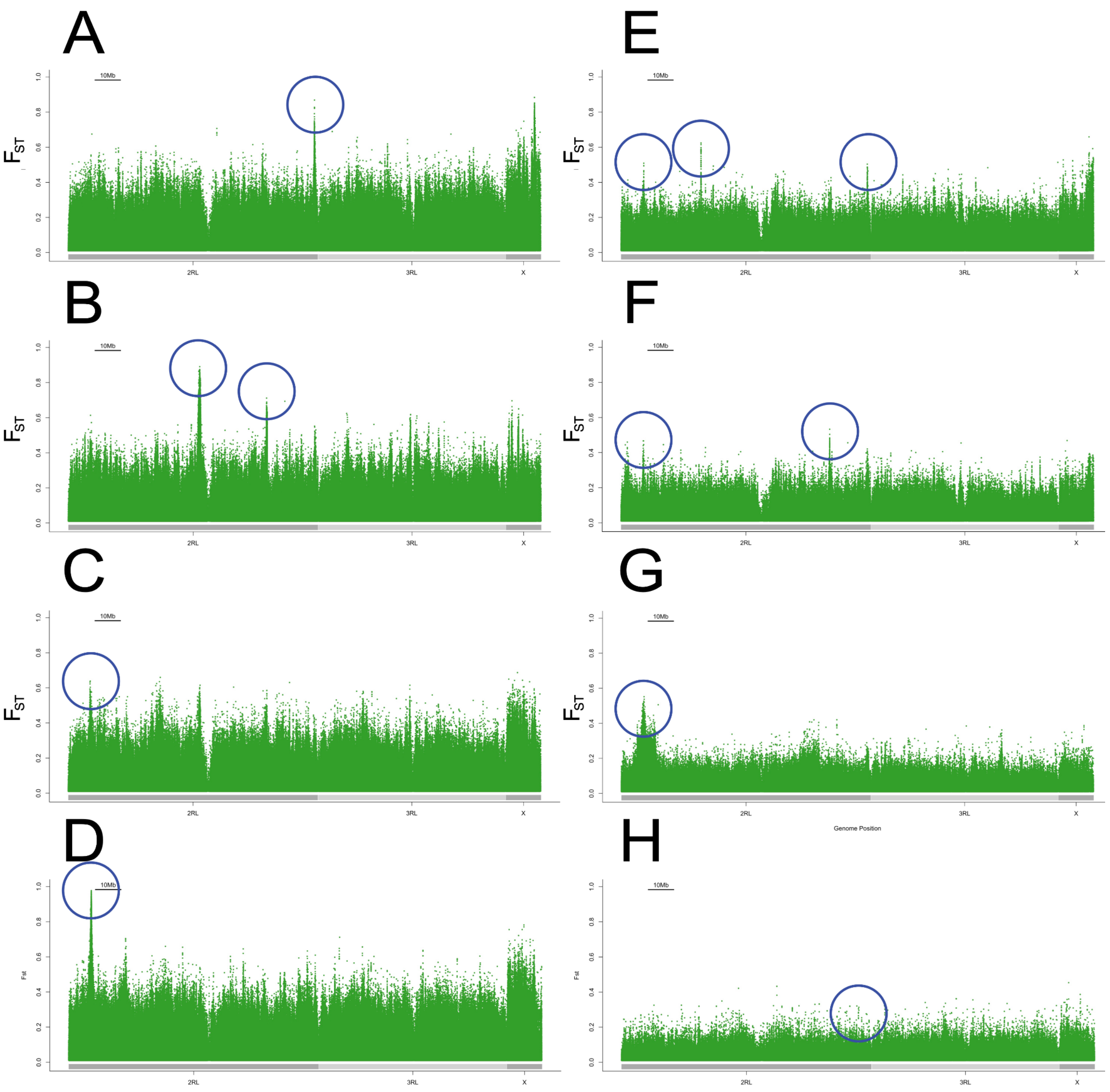


**Fig. S13**.

**Selection signals in genome-wide $F_{ST}$ scans.** Peaks highlighted in table S5 are circled, defined by sets of at least three autosomal variants with $F_{ST}$ greater than ten times the mean autosomal $F_{ST}$ for that comparison, spanning at least 500 bp, but with no more than 200 kb between adjacent variants. **(A)** $F_{ST}$ between Manaus (Amazonas, Brazil) and other Amazonas, Brazil sites. **(B)** $F_{ST}$ between Barcelos (Amazonas, Brazil) and other Amazonas, Brazil sites. **(C)** $F_{ST}$ between Coari (Amazonas, Brazil) and other Amazonas, Brazil sites. **(D)** $F_{ST}$ between Loreto (Peru) and all Amazonas, Brazil sites. **(E)** $F_{ST}$ between Blondin (French Guiana) and other French Guiana sites. **(F)** $F_{ST}$ between Cacao (French Guiana) and other French Guiana sites. **(G)** $F_{ST}$ between Guainía (Columbia) and Atures (Venezuela). **(H)** $F_{ST}$ between San Francisco de Tauchigadó (Chocó, Colombia) and Tagachí (Chocó, Colombia).

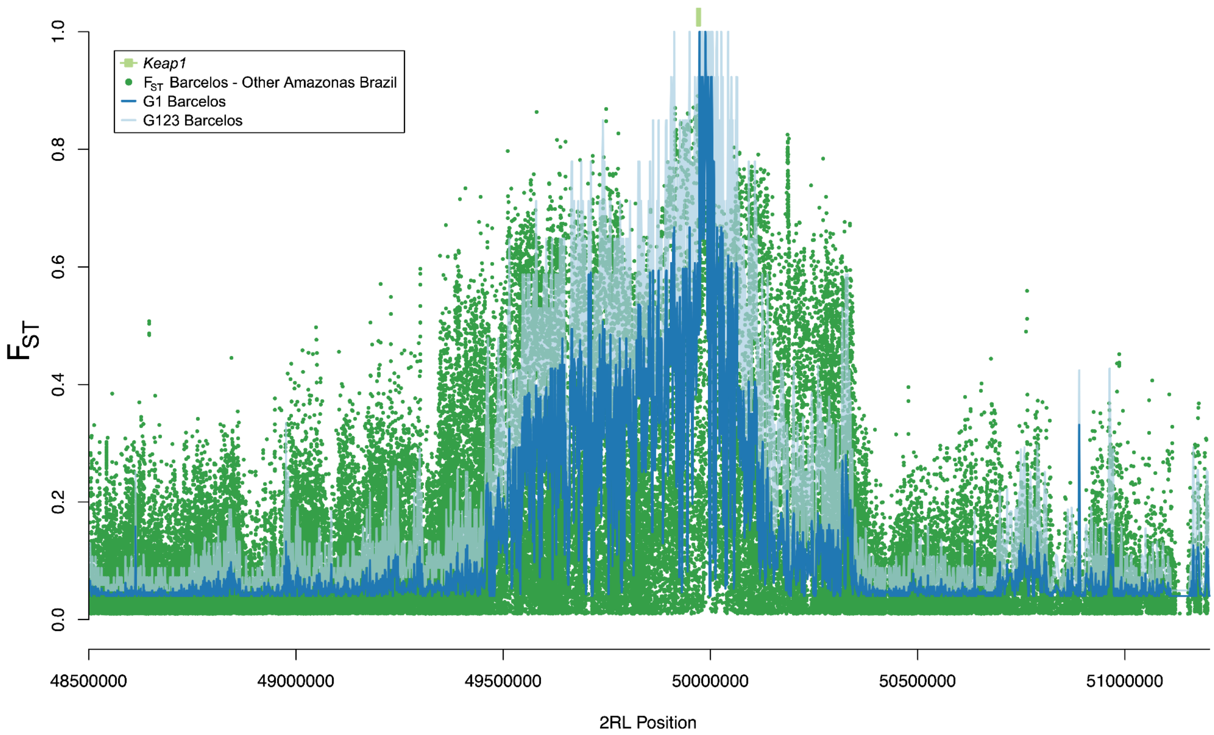


**Fig. S14**.

**Selection signal at 2RL_49.950-50.050 Mb in Barcelos, Amazonas.** Other than the P450 region at 2RL_8.425-8.575 Mb (Fig. 4), this is the strongest convergence of $F_{ST}$ outliers (table S5, fig. S13B) with G1/G123 outliers (tables S6 and S7). The top outlier gene for $F_{ST}$, which also overlaps the peak of G1 and G123, is *ADAR2_006983* (*Keap1*), known to play a role in metabolic resistance to insecticides.

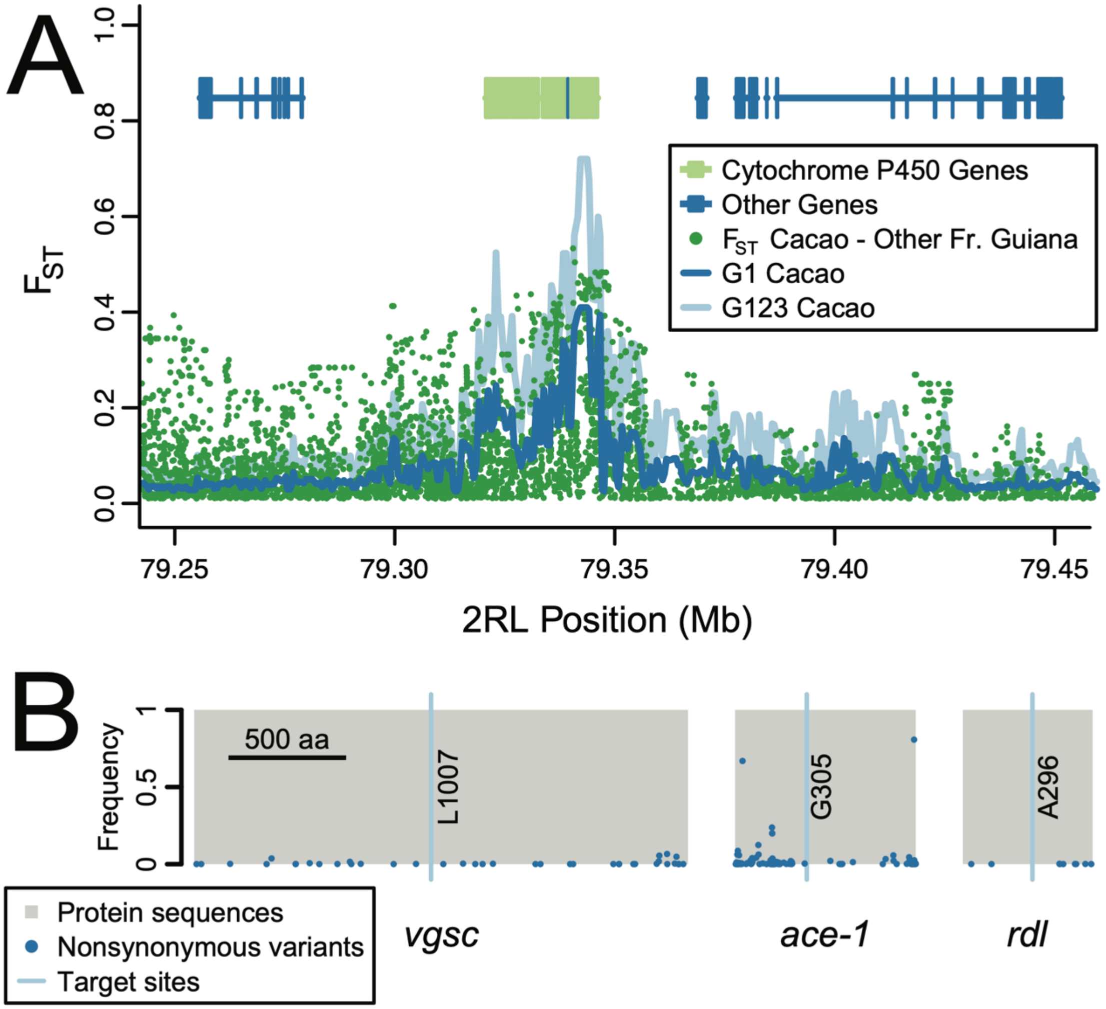


**Fig. S15**.
**Polymorphism at other potential insecticide resistance genes. (A)** Positive selection on a P450 gene cluster in Cacao, French Guiana, showing the highest $F_{ST}$ (dark green) in the genome when compared to the French Guiana populations, and also peaks at G1 (dark blue) and G123 (light blue). **(B)** No nonsynonymous polymorphism at known target site codons (light blue lines) in insecticide resistance genes (grey boxes): *vgsc*, *ace-1*, and *rdl*, and few common nonsynonymous variants elsewhere in the genes (dark blue dots, plotted as non-reference allele frequency).
